# Spatiotemporal variations of surface water microplastics near Kyushu, Japan: A quali-quantitative analysis

**DOI:** 10.1101/2021.03.22.436354

**Authors:** Tsunefumi Kobayashi, Mitsuharu Yagi, Toshiya Kawaguchi, Toshiro Hata, Kenichi Shimizu

## Abstract

Microplastics in the ocean are threatening marine ecosystems. Although plastic contaminants are ubiquitous from rivers to polar oceans, their distribution is thought to be heterogeneous, implying that both spatial and temporal variability exist. Here, we elucidate the significant spatial and temporal (seasonal) variations in the quanti-qualitative characteristics of microplastics off the west coast of Kyushu, Japan in the East China Sea. Six surveys across nine stations (n = 54) were conducted over a 14-month period, and a total of 6131 plastic items were identified. The average microplastic abundance (items ·m^-3^) and size (mm) ± S.D. were 0.49 ± 0.92 (n = 54), and 1.71 ± 0.93 (n = 6131), respectively. Differences between the highest and lowest abundances were 50-fold among monthly means (1.97 ± 1.49, n = 9; 0.04 ± 0.03, n = 9), and 550-fold across all net tows (5.50; 0.01). With respect to colour, polymer type, and shape, white and transparent (68.5%), polyethylene (80%) fragments (76.0%) were the dominant composition. There were statistically significant differences for each of the analytical microplastic parameters among survey months (p < 0.02). Our results provide baseline data, and lead to a more comprehensive understanding of the spatiotemporal characteristics of microplastic pollution.

**Highlights:**

- Significant spatiotemporal variability in microplastic litter was detected based on the quali-quantitative analyses.
- Average (± S.D.) microplastic abundance was 0.49 ± 0.92 (items ·m^-3^) and size was 1.71 ± 0.93 (mm).
- Differences between highest and lowest abundances were 50-fold among monthly means, and 550-fold across all net tows.
- White and transparent polyethylene fragments were the dominant composition.

## 1. Introduction

Oceanic plastic pollution is a global-scale issue. Plastic products have been an indispensable part of modern society since the 1940s and 50s due to their favourable properties, such as low cost, light weight, high durability, and ease of design (Cole et al., 2011; Hammer et al., 2012). Approximately 80% of all manufactured, mass-produced plastics to date, however, have accumulated in landfills or natural environments (Geyer et al., 2017), and inappropriate management of waste plastics can lead to their transfer to marine environments (Gregory, 2009; Ryan et al., 2009; Kuroda et al., 2020). Approximately 10% of the produced plastics accumulate and persist in the marine ecosystems (Thompson, 2006), with an estimated 1.15–2.41 million tons introduced into the ocean via the river system every year (Jambeck et al., 2015; Lebreton et al., 2017). If current waste management trajectories are maintained, an additional ∼12,000 million metric tons of plastic waste will be deposited into landfills or the natural environment by 2050 (Geyer et al., 2017), and the amount of plastic in the ocean will surpass that of fish by weight (World Economic Forum 2016).

Plastic debris in the ocean is deteriorated by ultraviolet rays and waves, and when it degrades to a size of ≤ 5 mm, it is commonly referred to as microplastic (MP; Andrady, 2011). MPs are ubiquitous in the global oceans, and readily accumulate in marine ecosystems due to their small size and ease of ingestion by various marine organisms (De Witte et al., 2014; Auta et al., 2017). Once ingested, toxic substances amass inside an organism and lead to intestinal blockage or physical damage (Jovanovic, 2017); therefore, negative impacts on marine organisms can be precluded by concentrating on MP management.

Drifting MP on the sea surface are heterogeneously distributed throughout the globe, and positively correlate with epicentres of dumping in the Southeast Asian countries, including China (Jambeck et al., 2015). Accordingly, Isobe et al. (2015) reported total MP particle concentrations in the East Asian seas around Japan that were 16 and 27 times greater than those of the North Pacific Ocean and the world’s oceans, respectively. Furthermore, marine MPs can vary with the seasons or currents (Moore, 2008; Martinez et al., 2009; Doyle at al., 2011); hence, studies with inadequate transect numbers and/or over short time scales fail to capture the spatiotemporal patterning in abundance and characteristics (Ryan et al., 2009; Cole et al., 2011). Information on the long-term quantitative (abundance and size) and qualitative (shape, colour, and polymer) characteristics of MPs is limited, even though such studies would provide valuable patterning and baseline data for all future research (Cole et al., 2011). Additionally, data on MPs in the waters off Kyushu, Japan, which is a purported highly polluted area, are lacking. It was hypothesised in the study here that the significant spatiotemporal changes in abundance and type of sea surface MPs could be detected by long-term surveys; thus, repeated sea surface surveys over 14 months in the East China Sea were conducted to determine abundance, size, and type of MPs, and obtain the first regional assessment of spatiotemporal variations for this contaminant.

## 2. Materials and methods

### 2.1. Sampling design

Six sea surface surveys were carried out from April 2019 to June 2020 using training vessel *T/V* Kakuyo-maru (155 gross tonnage: Faculty of Fisheries, Nagasaki University), with each survey containing nine sampling stations (Fig. 1; Supplementary Table S1). A total of 54 samples were collected with neuston net (JMA, RIGO Co., Ltd., Tokyo, Japan; rectangular mouth opening, 0.75 m × 0.75 m; length, 3 m; mesh size, 350 μm) originally designed for sampling neustonic organisms such as zooplankton, fish larvae, and eggs near the sea surface (Isobe et al., 2015). According to the net mesh size, MPs ranging from 350 μm to 5 mm were targeted. To measure the volume of water that passes through during sampling, a flow meter (5571A, RIGO Co., Ltd., Tokyo, Japan) was attached to the net mouth. The net was towed from a boom installed at the bow-port side of the vessel to ensure that it was kept 5 m from the ship’s side and prevent debris disturbance by the bow wave. The net was towed around each station for ∼10 minutes at a vessel log speed of 2.0 knots (Doyle et al., 2011; GESAMP, 2019). Significant wave height and wind speed during sampling were also recorded by the onboard monitoring system (Supplementary Figs. S1). After sample retrieval, nets were rinsed with filtered sea water to ensure that all debris and organisms were attached to the cod end. Contents of the cod end were then carefully put into a sample bottle (2000 ml), and debris larger than 100 mm had their surface rinsed with filtered sea water in a 350 μm strainer net prior to removal. The samples were preserved with a 5% formalin solution buffered with sodium borate, and stored at room temperature until sorting.

**Figure 1.**
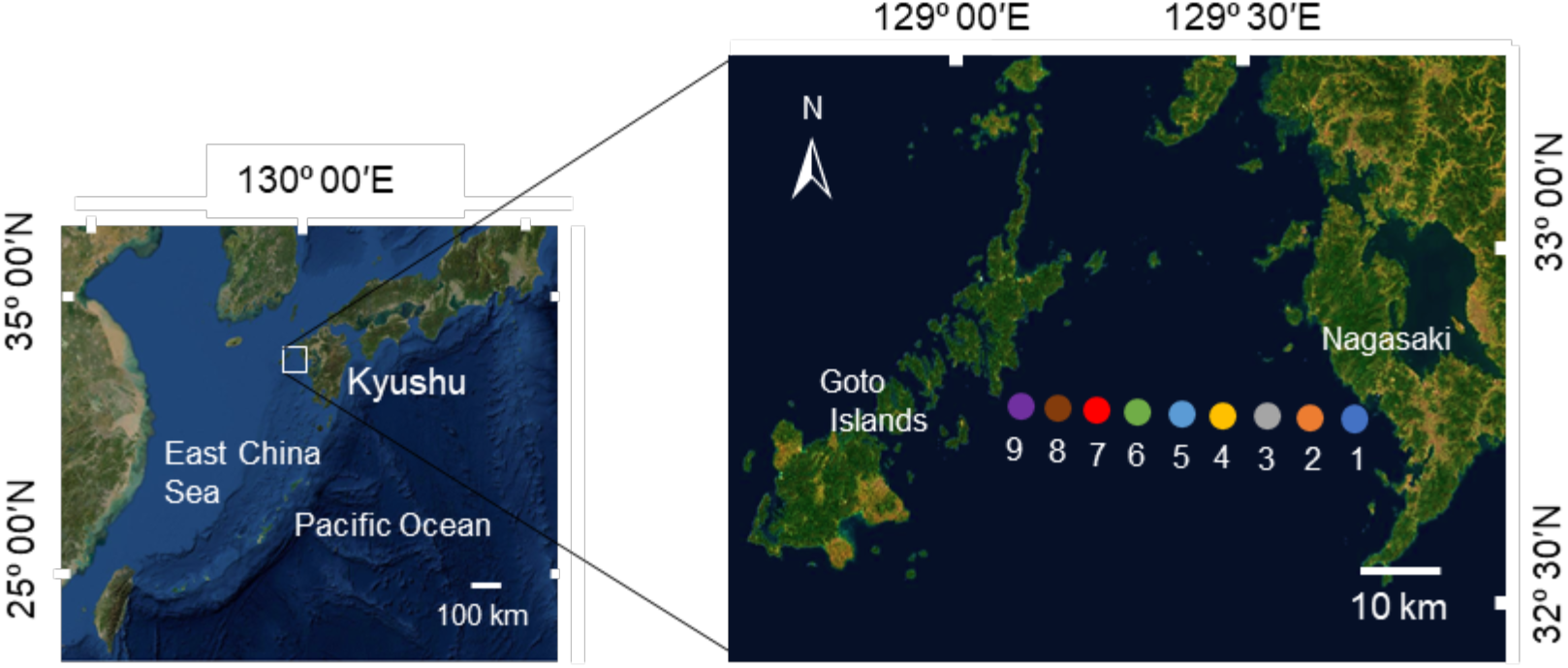
Sampling sites for sea surface microplastic collection in the water off the west coast of Kyushu, Japan. See Table S1 for the details of sampling locations, including coordinates, and water depth.

The volume of filtered water *V* (m^3^) was calculated from the flow meter readings according to Eqn (1):

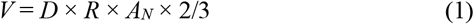

where *D* is a constant in the towing distance per revolution of the flow meter (0.107 m); *R* is the number of revolutions of the flow meter; *A*_*N*_ is the area of the net mouth (0.5625 m^2^); and 2/3 is the proportion of the neuston net frame that was submerged during towing, where the net was equipped with floats (buoys) that were placed to collect MPs from the upper ∼0.5 m surface layer. *D* represented the filtration efficiency that was calculated by towing the same distance with and without the net in a water tunnel at Tokyo University of Marine Science and Technology (Michida et al., 2019). Although areal units (e.g., items per km^2^) are frequently reported in similar research, when comparing wider areas such as the Mediterranean Sea and Atlantic Ocean (Collignon et al., 2012; Eriksen et al., 2013; Silvestrova and Stepanova, 2021), we adopted a volumetric approach (number of items per cubic meter of seawater) as the standardised measurement of debris for each sampling station; however we can still compare areas *A* (m^2^) by converting the recorded values according to Eqn (2):

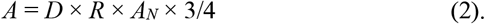

Furthermore, considering that the number of small plastic particles decreases exponentially with depth, it has been recommended that the concentration of MPs should be integrated vertically (Kukulka et al., 2012; Reisser et al., 2015). Theoretical concentration per square meter (items ·m^-2^) of MP particles on the surface can also be calculated by accounting for wind speed and significant wave height during survey (Supplementary Fig. S1).

### 2.1. Sample analyses

Sorting and identification of MPs were processed at the Fish and Ships Laboratory of the Faculty of Fisheries, Nagasaki University, using the methods described in Michida et al. (2019). To reduce sample contamination before and during analysis, the laboratory was closed to minimise airflow by closing windows and door. All equipment and working counters were disinfected with 90% alcohol solution prior to analysis, and experimental equipment was rinsed with deionised water prior to analysis. Equipment was covered with aluminium foil while not in operation. Additionally, during sorting, petri dishes with distilled water were placed within the work area and subsequently observed under a microscope to check for airborne contamination (Lam et al., 2020). Samples were visually examined by dissecting stereo microscopy (SMZ745T, Nikon Corporation, Tokyo, Japan), and debris was initially categorized as plastic and non-plastic by differentiating their colour, shape, and physical response properties (e.g., softness and texture) (Doyle at al., 2011; Hidalgo-Ruz et al., 2012). Non-plastic debris consisted of zooplankton, fish larvae and juveniles, crustaceans, wood fragments, and algae. In instances where many non-plastic particles were identified and difficult to sort, samples were treated with 30% H_2_O_2_ at room temperature for one week to dissolve interfering organic matter (Michida et al., 2019), sorted, and individually stored in pill cases.

All particles were photographed by a digital camera (NOA630, WRAYCAM Corporation, Tokyo, Japan) attached to a microscope, and the longest side of the major axis was measured using the *MicroStudio* software. MPs were categorized by four shape types (fragment, form, film, fibre) and eight colours (white, transparent, black, yellow, green, blue, red) based on the methods of Cheung et al. (2016) and Lam et al. (2020). Finally, all plastic polymer compositions were identified using an attenuated total reflection Fourier transform infrared spectrometer (ATR-FTIR; FT-IR-4600, JASCO Corporation, Tokyo, Japan) that exhibited a spectrum range from 4000–400 cm^-1^ at a resolution of 4 cm^-1^ and performed eight scans per sample. Background blank scans per every 50 samples were conducted for comparison, and the top-plate and diamond prism were cleaned with 75% ethanol prior to each sample analysis. The polymer type was identified by comparing a standard reference from KnowItAll Spectroscopy Library (Wiley Science Solutions, New Jersey, USA), and *PEAKPICK* function was adopted to confirm ≥ three plastic polymer characteristic peaks in the spectrum. The Hit Quality Index (HQI) was also recorded.

### 2.3. Statistical analysis

Concentration values were expressed in terms of the number of plastic items per cubic meter of seawater (items ·m^-3^), and all statistical data were expressed as mean items ± 1 standard deviation (S.D.). Since the quantitative data posed a non-nominal distribution (Shapiro-Wilk test: abundance, p = 1.1 × 10^−11^; size, p = 1.1 × 10^−4^) and the variance exhibited hetero-homogeneity (Bartlett test: abundance, p = 2.2 × 10^−16^, size, p = 6.8 × 10^−6^), a non-parametric analysis (Kruskal-Wallis test) was used. If the test indicated a significant difference, multiple comparisons were then performed using a Steel-Dwass post-hoc test. To test the differences among qualitative data (i.e., colour, shape, and polymer composition), Chi-square (χ^2^) tests were applied. All statistical analyses were performed in *R* (*v*.1.2.5033).

## 3. Result

### 3.1. Quantitative characterization of MP items

A total of 6131 plastic items were identified in this survey. The mean and median abundance were 0.49 ± 0.92 items ·m^-3^ (n = 54, ± S.D.), and 0.12 ± 0.56 items ·m^-3^ (n = 54, ± median absolute deviation—M.A.D.), respectively (Table 1). The Kruskal-Wallis test showed a significant difference in the mean abundance between months (n = 6, p < 0.0001; Fig. 2), while that observed among the sampling stations was insignificant (n = 9, p = 0.95). The highest observed abundance between surveys was 1.97 ± 1.49 items ·m^-3^ (October 2019), which was ∼50-fold higher than the lowest value: 0.04 ± 0.03 items·m^-3^ (April 2019; Fig. 2). Furthermore, the highest abundance among the sampling stations was 5.50 items·m^-3^ (October 2019), which was 550-fold higher than the lowest value: 0.01 items·m^-3^ (April 2019).

**Table 1.** Microplastic abundance: number of plastic items per cubic meter of seawater (items·m^-3^), and longest size of major axis (mm) collected from the surface waters off the west coast Kyushu, Japan. S.D.: standard deviation, and M.A.D.: median absolute deviation.

| Month | N | Abundance (items·m<sup>-3</sup>) |  | N | Size (mm) |  |
| --- | --- | --- | --- | --- | --- | --- |
|  |  | Mean ± S.D. | Median ± M.A.D. |  | Mean ± S.D. | Median ± M.A.D. |
| Apr. 2019 | 9 | 0.04 ± 0.03 | 0.02 ± 0.03 | 40 | 2.32 ± 1.05 | 2.00 ± 0.80 |
| Jun. 2019 | 9 | 0.09 ± 0.08 | 0.07 ± 0.06 | 85 | 1.96 ± 0.63 | 1.96 ± 0.48 |
| Jul. 2019 | 9 | 0.27 ± 0.31 | 0.18 ± 0.23 | 563 | 2.10 ± 0.32 | 2.22 ± 0.23 |
| Oct. 2019 | 9 | 1.97 ± 1.49 | 1.65 ± 1.00 | 4343 | 1.61 ± 0.19 | 1.66 ± 0.17 |
| Nov. 2019 | 9 | 0.29 ± 0.24 | 0.17 ± 0.20 | 608 | 2.13 ± 0.25 | 2.21 ± 0.21 |
| Jun. 2020 | 9 | 0.25 ± 0.45 | 0.06 ± 0.28 | 492 | 1.81 ± 0.36 | 1.84 ± 0.21 |
| Total | 54 | 0.49 ± 0.92 | 0.12 ± 0.56 | 6131 | 1.99 ± 0.57 | 1.93 ± 0.39 |

**Figure 2.**
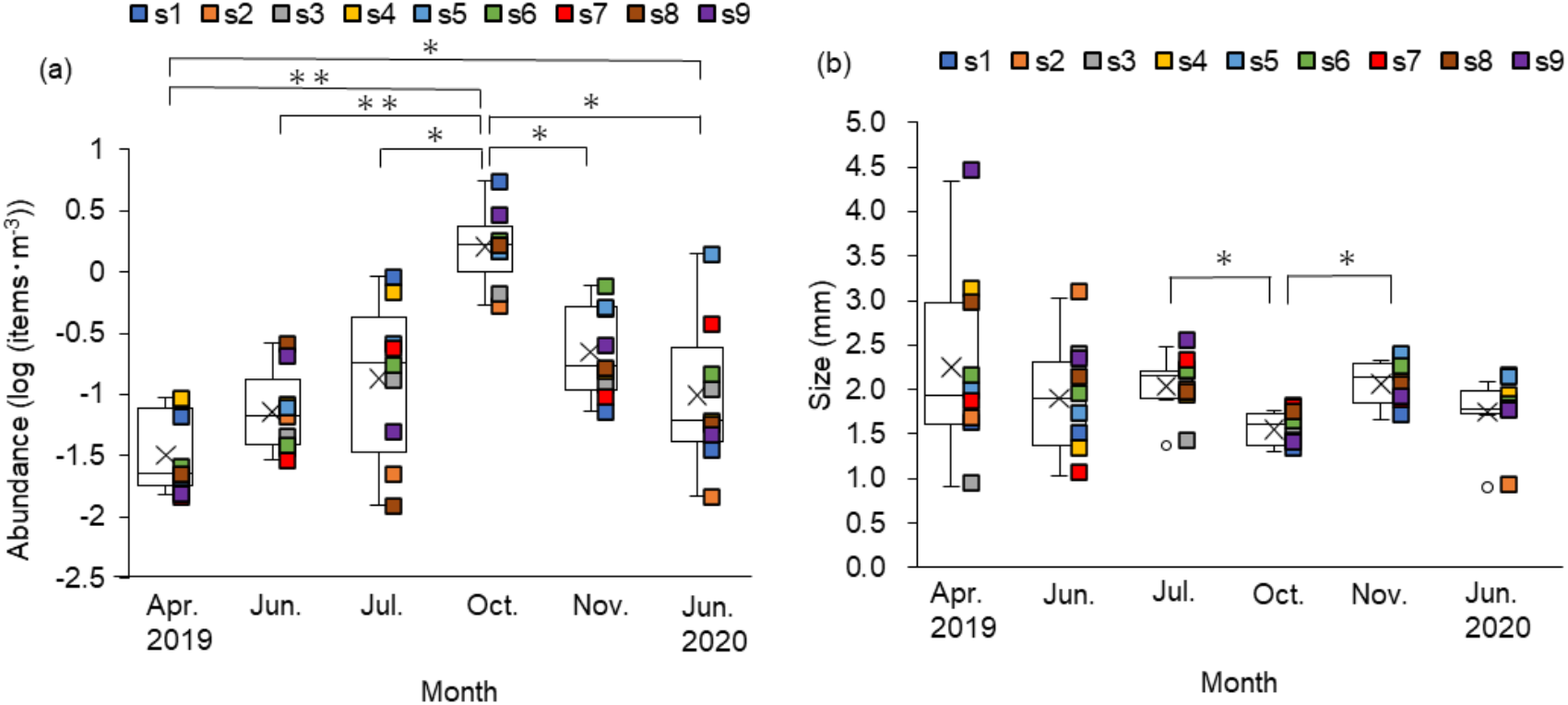
Box-and-whisker plots of the spatiotemporal patterning of (a) abundance, and (b) longest size on the major axis of microplastics in the waters off the west coast of Kyushu, Japan (* p < 0.05 and ** p < 0.01 were determined by the Steel-Dwass post-hoc test). The y-axis in (a) is in log10 scale. Lower and upper box boundaries indicate the 25^th^ and 75^th^ percentiles, cross marks represent the mean, interior box lines are the median, and exterior whiskers are the 10^th^ and 90^th^ percentiles. Plot abundance at each sampling station was n = 6.

The mean and median size of MPs were 1.71 ± 0.93 mm (n = 6131) and 1.45 ± 0.73 mm (n = 6131; Table 1), respectively. The Kruskal-Wallis test confirmed a significant difference in the mean sizes among the months (n = 6, p < 0.05; Fig. 3). Combining all months and stations, there was a trend of increasing abundance in MPs with decreasing mean size when x ranged from 1–5 mm or when it was < 1 mm (Fig. 4a). These trends were also confirmed by the size frequency in each month (Figs. 4b–g).

**Figure 3.**
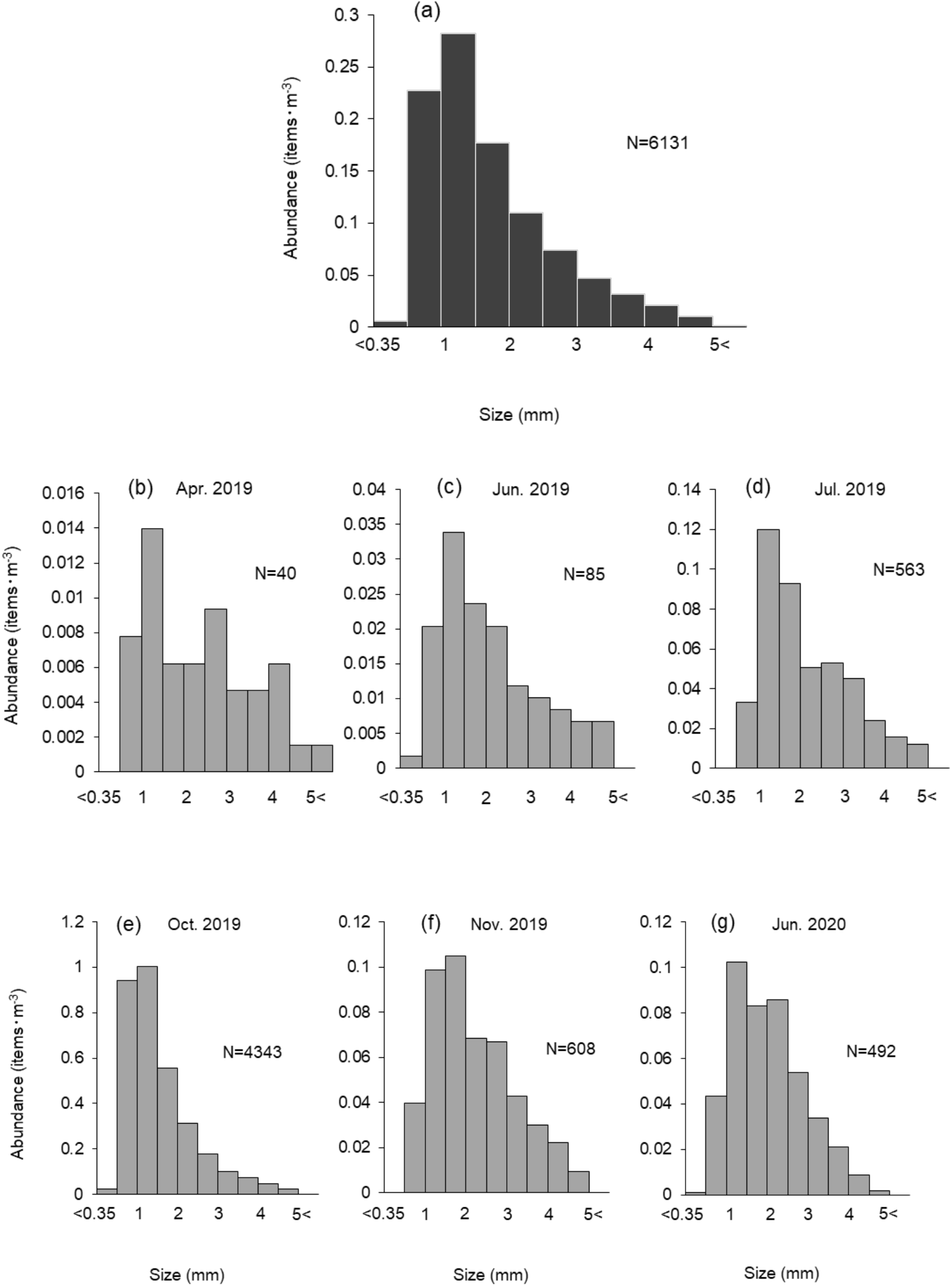
Size distribution of microplastic items in (a) total and (b)–(g) among each sampling date, showing mean abundances (items·m^-3^).

**Figure 4.**
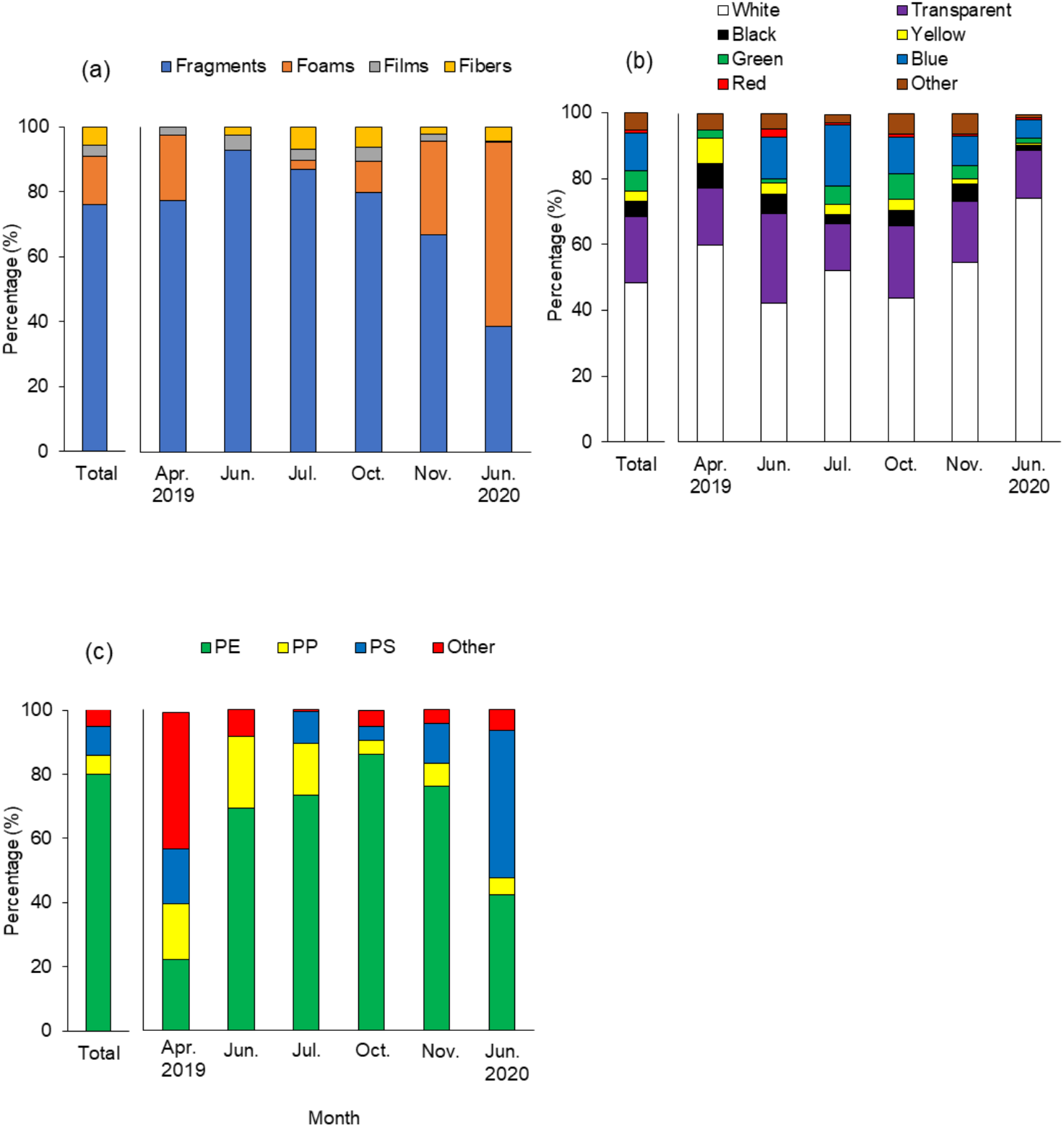
Total and monthly percentage compositions for (a) shape (fibres, films, foams, and fragments), (b) colour (white, transparent, black, yellow, green, blue, red, and other), and (c) polymer type (polyethylene, PE; polypropylene, PP; polystyrene, PS) and other microplastic abundances collected from the surface waters off the west coast Kyushu, Japan.

### 3.2. Qualitative characterization of MP items

Combining all months and stations, fragments were the primary shape type (76.0%), followed by foams (14.8%), fibres (5.6%), and films (3.5%); meanwhile, no notable primary MP such as microbead and resin pellet were found (Fig. 5a). Furthermore, white was the most abundant colour (48.3%), followed by transparent (20.2%) and blue (11.4%; Fig. 5b). With regards to polymer composition, polyethylene (PE) was the predominant type (80.0%), followed by polystyrene (PS; 9.0%) and polypropylene (PP; 5.9%; Fig. 5c). Other polymer types identified included polyvinyl chloride (PVC; 0.9%) and polyethylene terephthalate (PET; 0.4%). Although overall trends of qualitative characterization were roughly upheld in each individual month, Chi-square (χ^2^) tests indicated significant differences in shape, colour, and polymer type between them (p < 0.0001 for each qualitative variable).

## 4. Discussion

The work presented here represents the first report on a stationary, long-term survey of sea surface MPs from the marine waters surrounding Japan. Results from this study revealed that sea surface MPs displayed statistically significant spatiotemporal variability of quantitative (abundance and size) and qualitative (shape, colour and polymer type) characteristics in the waters off the west coast of Kyushu, Japan. These results strongly supported the hypothesis of heterogeneous MP distribution by space and time (Doyle et al., 2011; Gajšt et al., 2016; Lam et al., 2020). Indeed, differences between the highest and lowest average monthly abundances were 50-fold and 550-fold between all net tows, respectively.

In the study area, MP items were found in each of the 54 tows used for sampling over the 14-month analysis period (six surveys x nine stations). Although spatiotemporal heterogeneity of MP abundance was observed in the study area, there was no significant difference among sampling stations during survey period, thereby suggesting that surface MPs do not locally aggregate at a particular station within the survey area. Sea surface MPs can be found globally from semi-closed bays (Lattin et al., 2004; Chen et al., 2018; Kashiwabara et al., 2021; Nakano et al., 2021) to the Arctic Ocean (Peeken et al., 2018; Kanhai et al., 2020). Mean abundance of MPs are observed to be ten orders of magnitude, ranging from 10^−5^ items·m^-3^ in the equatorial Pacific (Spear et al., 1995) to 10^5^ items·m^-3^ off the southern coast of Korea (Song et al., 2014), depending on sea area and mesh size of the net. Mesh size of the neuston or manta nets primarily affects the absolute amount of plastic particles (Kang et al., 2015; Barrows et al., 2017; Lindeque et al., 2020; Tokai et al., 2021). Globally, mean abundance in the surface waters collected by a net with 290–350 µm mesh size was 0.96 ± 2.05 items·m^-3^ (Shim et al., 2018). In the present study, the average mean abundance was 0.49 ± 0.92 items·m^-3^, as evaluated by a neuston net with a 350 µm mesh size; this value was approximately half of the global average mean abundance. This value was similar to that found in the Tokyo Bay in May 2019 using the same neuston net (0.15–0.90 items·m^-3^, mean = 0.53, n = 4), except for one outliner (Nakano et al., 2021). Contrarily, Isobe et al. (2015) reported an abundance in the eastern Asian Seas of 3.74 items·m^-3^, and deemed the area a hotspot of sea surface MPs. The reasons for this discrepancy are unknown, but it may be related to the spatial and/or temporal variability of MP abundance seen in the present study. In fact, the mean abundance recorded in Tokyo Bay including the outliner in May 2019 (17.75 items·m^-3^) shot up to 3.98 items·m^-3^, leading the authors to conclude that the bay is one of the most polluted areas in the world with respect to MPs (Nakano et al., 2021). Our study also found considerable variability in abundance, ranging from 0.01 to 5.50 items·m^-3^; thus, further multiple samplings with repeated surveys are needed to better assess and characterize variations in abundance and distribution patterns.

The observed variability in mean abundance may not be caused by physical disturbances such as wave height, wind velocity, or close-range transport. Buoyant MP particles are mixed vertically and distributed within the upper water column by wind and turbulent transport; thus, surface abundance decreases with increasing wind velocity (Kukulka et al., 2012; Reisser et al., 2015). Although wind-driven mixing was likely a factor throughout the present study, this physical turbulence could not explain the observed variability alone as wind velocity during sampling did not correlate with either abundance or a significant wave height (Fig. S1). Furthermore, nearby transport arising from adjacent rivers and landfills also could not explain the observed variability in abundance. The high unidirectional flow of rivers drives the movement of plastic particles into the ocean (Browne at al., 2010; Moore et al., 2002). During rainy seasons, surface runoff increases and releases MPs from inland areas to streams and rivers (Williams and Simmons, 1999; Cunningham and Wilson, 2003; Nakano et al., 2021), resulting in increased MP concentrations of the adjacent coastal sea areas (Cheung et al., 2016); however, the abundance of MP items in the survey area of the present study did not increase during the rainy season. Therefore, variability in abundance here may have arisen from long-range transport, rather than short. The hydrodynamic environment of this study area is strongly influenced by the Kuroshio current (Katoh et al., 1996). Isobe et al. (2015) showed that small plastic fragments near Japan (upstream from the East and Southeast Asian countries) were responsible for discharging large amounts of plastic waste into the ocean (Jambeck et al., 2015) and occurred due to the Kuroshio and Tsushima currents. A northward branched channel from the Kuroshio currents was observed by a general ocean circulation model (Fig. S2); however, no clear relationship was found between geostrophic mean flow and the dramatic increase in abundance in October 2019 (Figs. 2 and S2). This suggests that the strength of northward branched currents in the study area could not explain the observed increase in abundance, and perhaps this explosion originated from massive clusters generated by extreme flash flooding. One storm resulted in litter being deposited at even greater distances from the river mouth and coastal area (Lattin et al., 2004). Particle tracing model analyses are needed to pinpoint the emission sources, and fixed-point observations are required to ascertain the future ecological consequences of these anthropogenic particles.

In this study, heterogeneity of MP characteristics was accompanied with measurements of mean size (Fig. 2b). Although many recent studies have reported MP sizes (e.g., Cózar et al., 2014; Reisser et al., 2015; Gajšt et al., 2016; Lam et al., 2020), information on seasonal variability of sizes are still lacking. Mean sizes in October 2019 were significantly smaller than those of the preceding months (July and November 2019; Fig. 2b). Plastic debris is gradually degraded as it moves through the ocean currents and gets repeatedly washed ashore on beaches, before returning to the ocean (Andrady 2011; Hidalgo-Ruz et al., 2012; Isobe et al., 2015). Thus, the observed difference in mean size could be due to more intense physical processes, longer drifting and/or residence times (i.e., older debris), or secondary plastics originating from larger debris. Furthermore, size distribution of MP items should be noted. Overall, the mode of item size was ∼1-1.5 mm and abundance decreased rapidly at sizes < 1 mm (Fig. 3a); these trends were also observed in every monthly size distribution (Figs. 3b–g). Mesh selectively was a major reason for the observed skewed distribution, as most MPs < 1 mm could not be collected by the mesh size used (350 µm; Tokai et al., 2021). Other factors, such as coastal deposition (Hinata et al., 2017), sessile organisms (Zettler et al., 2013; Long et al., 2015), and bioaccumulation by zooplankton (Cole et al., 2015; Desforges et al., 2015) also could have contributed to the patterns observed, as they play potentially significant roles in the removal of MPs from the surface waters.

Significant differences in qualitative characteristics (shape, colour, and polymer type) were also observed in the present study (Fig. 4). All MPs identified here were classified by secondary plastics. Fragments were the most frequent shape type observed (Fig. 4a); meanwhile, white and transparent colour types were predominant and later identified as PE by FT-IR analysis (Figs. 4b and c). These compositions are in agreement with previous studies carried out in the Pearl River Estuary (Lam et al., 2020), Tokyo Bay (Nakano et al., 2021), and the western North Pacific Ocean (Yamashita et al., 2007); however, the composition of all three characteristics varied significantly among survey months. For example, foam shapes were relatively high in June 2020 and accompanied by an increase in low density PS (polystyrene). The density of plastics varies with the polymer type (Browne et al., 2010). The density, along with the MP size and geographic emission variability, seem to largely affect transportation routes and the resulting spatiotemporal heterogeneity in MPs. More importantly, the qualitative and quantitative characteristics of MPs affect ingestive behaviour by marine biota. Ory et al. (2017) reported that fish selectively ingested specific colours of MPs more closely resembling zooplankton, a significant factor in the growing global concern of bioaccumulating MP particles found in the gastrointestinal tracts of fish (e.g., Collard et al., 2017; Lusher et al., 2015; Rummel et al., 2016). Once plastic waste is released into the ocean, complete retrieval is impossible, and there is urgent need for global cooperation before the amount or marine plastic will outweigh the fish.

## Credit authorship contribution statement

**Tsunefumi Kobayashi:** Formal analysis, Investigation, Writing - original draft, Visualization. **Toshiya Kawaguchi:** Investigation, Writing - review &editing. **Kenichi Shimizu:** Investigation, Writing - review &editing. **Toshiro Hata:** Investigation, Writing - review &editing. **Mitsuharu Yagi:** Conceptualization, Methodology, Investigation, Writing - original draft, Editing, Supervision, Funding acquisition.

## Acknowledgements

The authors are grateful to the captain, officers, and crews of the training ship *Kakuyo-maru*, and also acknowledge the “Fish and Ships laboratory” staff from the Faculty of Fisheries, Nagasaki University, who directly and indirectly helped throughout the entirety of the study. Finally, thanks to the editor and anonymous reviewers for their valuable comments and suggestions that greatly improved the quality of this manuscript.

