## Supplementary information for "Spatiotemporal variations of surface water microplastics near Kyushu, Japan: A quali-quantitative analysis"

Table S1. Sampling station number, coordinates, and depth.

| Sampling station | Latitude (N) | Longitude (E) | Depth (m) |
|---|---|---|---|
| 1 | 32º 46′ 29″ | 129º 43′ 14″ | 47 |
| 2 | 32º 46′ 36″ | 129º 39′ 27″ | 68 |
| 3 | 32º 46′ 39″ | 129º 34′ 33″ | 61 |
| 4 | 32º 46′ 46″ | 129º 29′ 14″ | 87 |
| 5 | 32º 46′ 56″ | 129º 24′ 10″ | 88 |
| 6 | 32º 46′ 57″ | 129º 18′ 16″ | 88 |
| 7 | 32º 47′ 12″ | 129º 13′ 05″ | 88 |
| 8 | 32º 47′ 16″ | 129º 07′ 49″ | 91 |
| 9 | 32º 47′ 28″ | 129º 03′ 09″ | 37 |

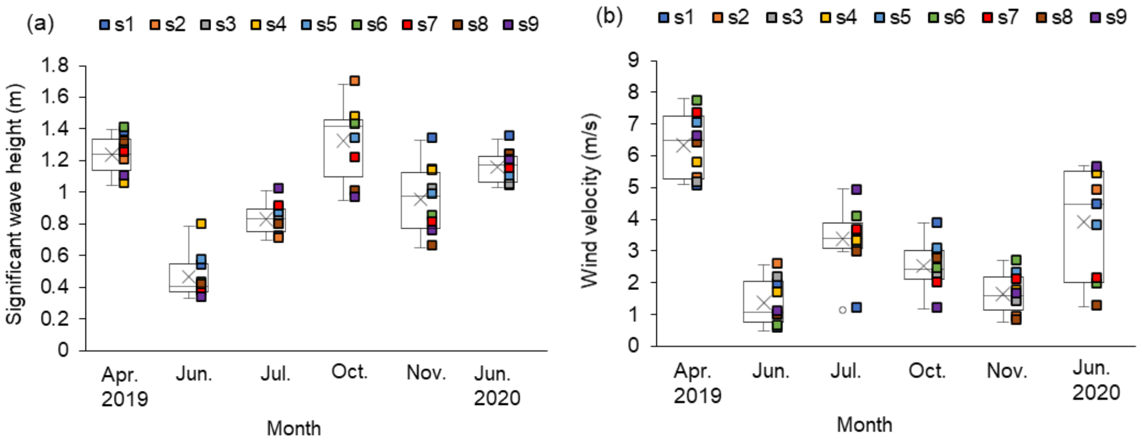


Figure S1. Box-and-whisker plots for: (a) significant wave height and (b) wind velocity during sampling. Upper and lower box boundaries are the 25$^{th}$ and 75$^{th}$ percentiles, respectively; cross marks are the mean; the internal line is the median; and the external whiskers are the 10th and 90$^{th}$ percentiles, respectively. n = 9 for each plot, corresponding to the number of sampling stations.

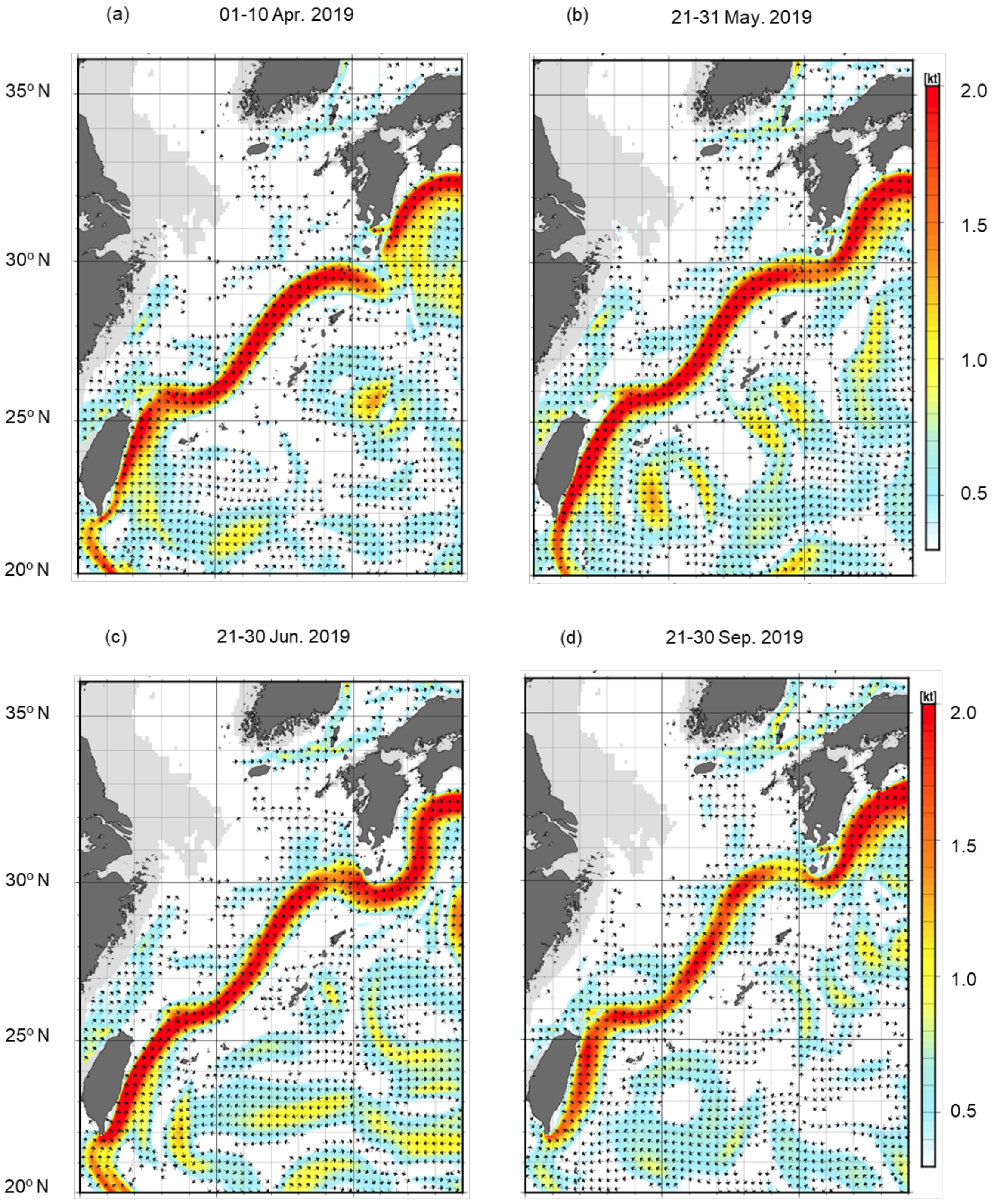
(a)
01-10 Apr. 2019
(b)
21-31 May. 2019
(c)
21-30 Jun. 2019
(d)
21-30 Sep. 2019
35° N
30° N
25° N
20° N
[kt]
2.0
1.5
1.0
0.5

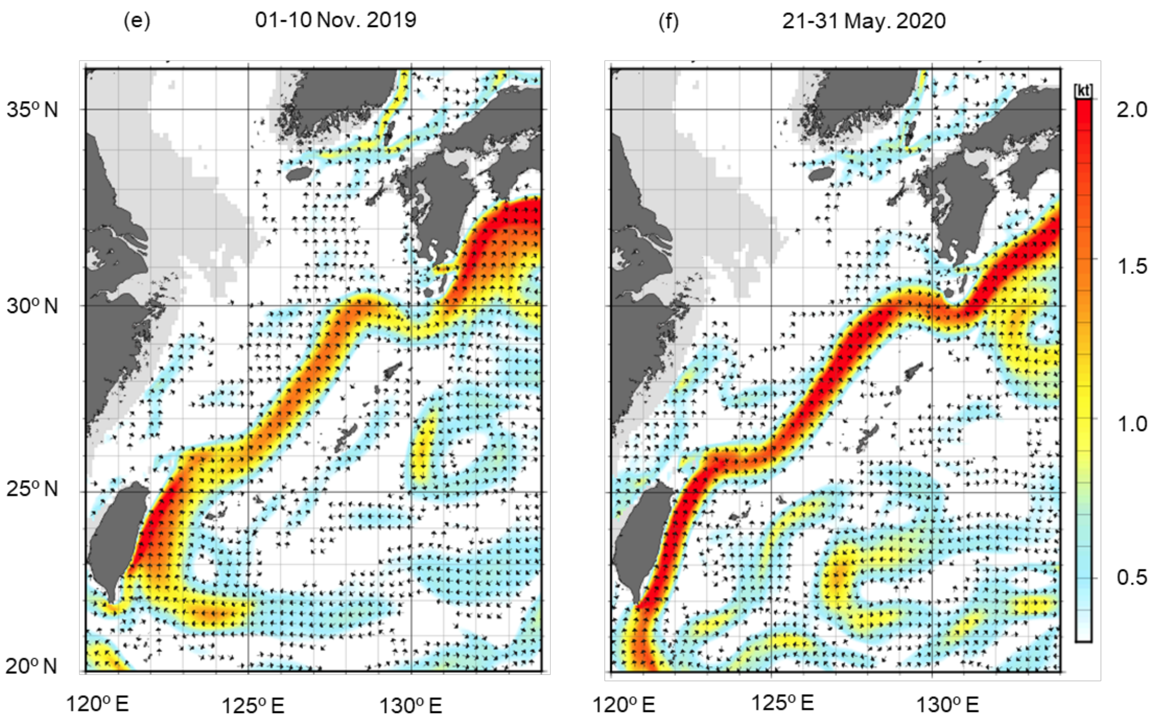


Figure S2. Ten day mean currents before each sampling date in the region (11 Apr. 2019, 3 Jun. 2019, 3 Jul. 2019, 4 Oct. 2019, 12 Nov. 2019, and 1 Jun. 2020). Current data were derived from an ocean general circulation model integrated with information from satellite, ship, meteorological float, and submerged buoy data, as produced and distributed by Japan Meteorological Agency (https://www.data.jma.go.jp/gmd/kaiyou/data/db/kaikyo/jun/current_HQ.html).
